# The genetic architecture of emerging fungicide resistance in populations of a global wheat pathogen

**DOI:** 10.1101/2020.03.26.010199

**Authors:** Danilo Pereira, Bruce A. McDonald, Daniel Croll

## Abstract

Containing fungal diseases often depends on the application of fungicidal compounds. Fungicides can rapidly lose effectiveness due to the rise of resistant individuals in populations. However, the lack of knowledge about resistance mutations beyond known target genes challenges investigations into pathways to resistance. We used whole-genome sequencing data and association mapping to reveal the multilocus genetic architecture of fungicide resistance in a global panel of 159 isolates of *Parastagonospora nodorum*, an important fungal pathogen of wheat. We found significant differences in azole resistance among global field populations. The populations evolved distinctive combinations of resistance alleles which can interact synergistically. We identified 34 significantly associated SNPs located in close proximity to genes associated with fungicide resistance in other fungi, including an MFS transporter. Using fungal colony growth rates and melanin production at different temperatures as fitness proxies, we found no evidence that resistance was constrained by genetic trade-offs. Our study demonstrates how genome-wide association studies of a global collection of pathogen strains can recapitulate the emergence of fungicide resistance. The distinct complement of resistance mutations found among populations illustrates how the evolutionary trajectory of fungicide adaptation can be complex and challenging to predict.

## Introduction

Fungal pathogens threaten global food security and human health (Fisher et al. 2012), causing economic losses and impacting global poverty (Strange & Scott 2005). Treatment of both animal and plant fungal infections rely on the application of fungicidal compounds that increasingly exhibit a decrease in effectiveness (Fisher et al. 2018). The emergence of resistance in fungal populations affects nearly all major fungicide groups (Stehmann & De Waard 1996; Sierotzki et al. 2000; Avenot & Michailides 2007). The loss in efficacy is due mainly to the intense selective pressure imposed by continuous fungicide applications based on single active compounds (Bosch & Gilligan 2008; Hobbelen et al. 2014). Mutations reducing fungicide sensitivity are strongly favored by selection and quickly increase in frequency in selected populations (McDonald & Linde 2002; Bosch & Gilligan 2008; van den Bosch et al. 2011). The genetic architecture associated with resistant phenotypes will arise from a complex array of mutations and their interactions, in turn affected by the pathogen population biology and characteristics of the fungicide. The mode of action (such as impairing mitochondrial respiration, Yang et al. 2011) and the number of target sites (single- vs. multisite fungicides) will play key roles in defining routes to resistance. Genetic trade-offs impacting fitness (Mikaberidze & McDonald 2015), innate resistance and epistatic effects will also significantly shape the evolutionary process of resistance emergence (Lucas et al. 2015). Hence, deciphering the genetic architecture of emerging fungicide resistance can provide useful insights and potentially identify key factors governing the evolutionary responses of pathogens.

Fungicides from the family of demethylation inhibitors (DMIs), in particular azoles, are the most widely used molecules in agriculture and human medicine (Fisher et al. 2018). Azoles hinder the biosynthesis of ergosterol through inhibition of the 14α-demethylase (CYP51) enzyme, negatively impacting the fungal cell membrane integrity and permeability (Georgopapadakou & Walsh 1996; Lass-Flörl 2011). In this group of fungicides, resistance emerges throughout different mechanisms, including (i) amino acid mutations in the target protein, (ii) overexpression of the gene encoding the target protein, and (iii) enhanced transporter activity reducing intracellular concentrations of the fungicide (Becher & Wirsel 2012; Cools & Fraaije 2013). Importantly, resistance in populations may be based on multiple mechanisms and is likely to be constrained by fitness costs (Zhan & McDonald 2013; Mikaberidze & McDonald 2015). Resistance can also emerge multiple times independently within species (Torriani et al. 2009). Structural changes in the CYP51 protein are considered the most common mechanism leading to resistance across species (Deng et al. 2007; Lucas et al. 2015). Highly resistant genotypes can acquire dozens of different mutations in the *CYP51* gene in a stepwise manner (Cools & Fraaije 2013). The consequences of the stepwise accumulation of mutations are complex interactions with the genetic background and selection for compensatory mutations (Cools et al. 2010; Lucas et al. 2015; Mullis et al. 2018). Alternative mechanisms to point mutations include copy-number variation of *CYP51* paralogs that are frequent in *Ascomycota* fungi (Brunner et al. 2015; Deng et al. 2007; Liu et al. 2011; Yan et al. 2011). The lack of knowledge about resistance mutations occurring outside of the *CYP51* gene limit our understanding of the likely importance of interactions among resistance mutations occurring in other genes. Genome-wide analyses of fungicide resistance will fill important gaps in our understanding of how resistance is acquired within species.

Knowledge of where resistance genes are located in the genome is needed to integrate information on standing genetic variation and evidence for recent selection. Genome-wide analyses led to the discovery of specific structural variation and single nucleotide polymorphisms (SNPs) underpinning fungicide resistance. A series of studies in *Candida albicans* established the contributions of variation in gene copy number (Selmecki et al. 2010), mutations in transcription factors (Coste et al. 2006; Dunkel et al. 2008), aneuploidy (Hill et al. 2013), and specific polymorphisms in over 240 genes (Ford et al. 2015) to fungicide resistance.

Population genomic analysis of 24 environmental and clinical strains of *Aspergillus fumigatus* revealed segregating azole-resistance alleles in different genetic backgrounds (Abdolrasouli et al. 2015). In the agricultural environment, the emergence of fungicide resistance is expected to be rapid (McDonald & Linde 2002; Croll & McDonald 2017) as a result of the genetic homogeneity of host plants and intensive fungicide usage (Stukenbrock & McDonald 2008). In addition to rare *de novo* mutations, the standing genetic variation from natural pathogen populations is a likely source for fungicide adaptation that is seldom explored (Barrett & Schluter 2008; Yamashita & Fraaije 2018). Very few studies have considered the genomic landscape of natural populations when investigating the evolution of fungicide resistance in agroecosystems (Mohd-Assaad et al. 2016; McDonald et al. 2019).

The haploid fungus *Parastagonospora nodorum* is a necrotrophic pathogen affecting wheat production worldwide (Oliver et al. 2012; Ficke et al. 2017). *P. nodorum* colonizes leaves and ears of wheat, causing necrotic lesions and reducing yield. *P. nodorum* spreads across regions on contaminated seeds and wheat straw (Solomon et al. 2006; Bennett et al. 2007). The main migration routes among China, Europe, North America and Australia were described in earlier studies (Stukenbrock et al. 2006). Most populations are characterized by frequent sexual recombination (Keller et al. 1997; Sommerhalder et al. 2006). A recent population survey in North America identified two major populations of *P. nodorum* with different genomic regions enriched in effectors under selection (Richards et al. 2019). This and other studies (Pereira et al. 2019) provide abundant evidence of the potential for *P. nodorum* to respond to selection in the agricultural environment. In Europe, most varieties of wheat are susceptible to *P. nodorum* (Downie et al. 2018), so high amounts of fungicide are applied to control this and other foliar diseases (Fones & Gurr 2015), impacting the entire cohort of wheat pathogens (Blixt et al. 2010; Knorr et al. 2019). European populations of *P. nodorum* were found to contain isolates with point mutations in major genes related to fungicide resistance including in *CYP51* (Blixt et al. 2009; Pereira et al. 2017). However, the genetic architecture associated with azole resistance emergence remains largely unexplored.

In this study, we analyze a collection of 159 *P. nodorum* genomes from seven field populations collected around the world and perform genome-wide association studies (GWAS) to establish the genetic basis of fungicide sensitivity globally. We also investigate whether the emergence of fungicide resistance led to pleiotropic effects using measures of fungal growth and melanization.

## Material and Methods

### Fungal populations

Isolates of *P. nodorum* were sampled from wheat <fields naturally infected by the pathogen. A total of 159 isolates chosen from seven fields (~20 isolates per field) were included in our analyses. The sampled countries/regions included Australia (2001), Iran (2005 and 2010), South Africa (1995), Switzerland (1999A and 1999B), New York (USA, 1991), Oregon (USA, 1993) and Texas (USA, 1992). All isolates were previously genotyped using microsatellite markers (Stukenbrock et al. 2006; McDonald et al. 2012). We selected only unique haplotypes for this study.

In earlier publications (Sommerhalder et al. 2006; Stukenbrock et al. 2006; McDonald et al. 2012, 2013; Pereira et al. 2017, 2019), the Switzerland 1999B population was indicated to originate from China in 2001. As a result of the genome sequence analyses reported in this paper, we believe that a transcription error led to mislabeling of the China 2001 population, which we now believe originated from a Swiss field of wheat located near Bern, ~150 km away from where the Swiss 1999A population was collected. The re-assignment of the China 2001 population to Switzerland 1999B does not compromise any of the analyses or interpretations reported in this manuscript.

### Fungicide sensitivity phenotyping

Isolates were recovered from long-term storage in silica gel at −80°C by placing silica gel fragments on the center of round Petri dishes containing potato dextrose agar (PDA, 4 g L^-1^ potato starch, 20 g L^-1^ dextrose, 15 g L^-1^ agar and 50 mg L^-1^ kanamycin). The plates were placed in chambers with a constant temperature of 24°C in the dark to induce mycelial development. After three days of growth, mycelium from each isolate was excised from the edges of the colonies with a cork borer (5 mm) and transferred to new PDA plates, to be used as the inoculum source for the sensitivity phenotyping experiment. All 159 isolates were phenotyped using four doses of propiconazole (Syngenta, Basel, Switzerland) chosen based on previous experiments that were conducted to determine the dose range that revealed the greatest variation in sensitivity among isolates. The selected doses were 0, 0.1, 0.5 and 1 ppm of propiconazole diluted in dimethyl sulfoxide (DMSO, 0.002% v/v). The doses of propiconazole or DMSO alone (as a control) were incorporated into molten PDA (~50°C) with a magnetic stirrer and a 50 ml volume was poured into square Petri dishes (120 x 120 x 17 mm, Huberlab).

Using a 5 mm cork borer, mycelial plugs were excised from the edge of colonies developing after seven days of growth on the inoculum plates. Four plugs from each isolate were placed in the corners of square plates with equidistant separation. Isolates were replicated twice, generating eight colonies in total for each of the four doses. Plates were randomized in an incubation chamber and grown at a constant temperature of 24°C and with no light during the entire experiment. Digital images of the colonies were acquired at 2, 4, 6 and 8 days after inoculation (DAI) through each plate lid. After image acquisition, plates were returned to the incubation chamber and randomized again. Images were analyzed using a batch script in ImageJ (Lendenmann et al. 2014; Schneider et al. 2012), yielding quantitative measures of each colony for total colony area (mm^2^) and melanization (mean grey values). The effective concentration that inhibited mycelial growth by 50% (EC_50_) was determined using a dose-response curve based on colony radius 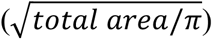 values in the R package *drc* version 3.0-1 (Ritz et al. 2015).

### Whole-genome sequencing

Fragments of mycelium from four-day-old colonies growing on PDA media were transferred to 50 ml Potato Dextrose Broth (PDB) media and cultured for 4 to 6 days at 24 °C while shaking at 120 rpm. The resulting mycelial colonies were filtered through sterile cheesecloth and lyophilized for 72h. Dried fungal material was used for DNA extraction with the DNeasy Plant Mini Kit (Qiagen) following the manufacturer’s standard protocol. We sequenced the genomes for all 159 isolates included in this study. The sequencing was performed on an Illumina HiSeq 2500 platform producing paired-end reads of 150 bp. Preparation of sequencing libraries and sequencing were performed at the Functional Genomics Center in Zurich. Raw sequence reads were deposited in the NCBI Short Read Archive under BioProject PRJNA606320.

### Genome alignment, variant calling and quality filtering

Raw reads were trimmed for remaining Illumina adaptors and read quality was assessed using Trimmomatic version 0.36 (Bolger et al. 2014) with the following parameters: illuminaclip = TruSeq3-PE.fa:2:30:10, leading = 10, trailing = 10, slidingwindow = 5:10, minlen = 50. Trimmed reads were aligned against the reference genome established for the isolate SN2000 (Richards et al. 2017). Reference genome mapping was performed using the short-read aligner Bowtie2 version 2.3.3 (Langmead & Salzberg 2012), using the –very-sensitive-local option. Picard tools version 2.17.2 was used to mark PCR duplicates (http://broadinstitute.github.io/picard). All sequence alignment (SAM) files were sorted and converted to binary (BAM) files using SAMtools version 1.2 (Li et al. 2009). Single nucleotide polymorphism (SNP) calling and variant filtration were performed using the Genome Analysis Toolkit (GATK) version 3.8-0 (McKenna et al. 2010). Initially, we used HaplotypeCaller on each isolate BAM file individually with the --emit-ref-confidence GVCF and -ploidy 1 options. Then, joint variant calls were produced using GenotypeGVCFs with the flag -maxAltAlleles 2. Finally, SelectVariants and VariantFiltration was used for hard filtering SNPs with the following cut-offs: QUAL < 200; QD < 10.0; MQ < 20.0; −2 > BaseQRankSum > 2; – 2 > MQRankSum > 2; −2 > ReadPosRankSum > 2. SNPs that failed the PASS designation by GATK were removed and we kept only bi-allelic sites. For the final dataset, we retained SNPs with a genotyping rate of at least 90% and a minimum allele frequency of 5% using vcftools version 0.1.15 (Danecek et al. 2011).

### Genome-wide association mapping

Association analysis was performed using the R package *GAPIT* version 2 (Tang et al. 2016), using a mixed linear method (MLM) (Yu et al. 2005). This model improves the control of false positives (type I errors) by incorporating fixed and random effects. Alternatively, we tested the inclusion of principal components (PCs) from a PC analysis to correct for population structure (Q) or a kinship matrix (K) to account for cryptic relationships (VanRaden 2008; Yu et al. 2005). We identified the most appropriate set of parameters and covariates by comparing the models MLM + K and MLM + K + Q, where Q stands for the three first PCs. Based on a Bayesian information criterion (BIC) (Schwarz 1978) analysis performed in *GAPIT*, the MLM + K model was selected as the most appropriate for our dataset (Table S1). We considered associations to be significant when *P* values were smaller than the Bonferroni threshold at a = 0.05 (*P* < 1.1 e-07). False discovery rate (FDR) thresholds of 5% (*P* < 7.15 e-07) and 10% (*P* < 8.26 e-06) were determined using the R package *q-value* version 2.18.0 (Storey & Tibshirani 2003). We explored the genomic regions containing significantly associated loci using bedtools version 2.29.0 (Quinlan & Hall 2010).

### Population structure and linkage disequilibrium analyses

Population structure was inferred using both a PC analysis in TASSEL version 5.2.56 and a model-based clustering implemented in STRUCTURE v2.3.4 (Pritchard et al. 2000; Bradbury et al. 2007). We visualized the two first PCs using the *ggplot2* package in R. The genetic markers used as input in STRUCTURE were composed of 2’348 SNPs. These SNPs were selected randomly across the genome using a sampling window of 10 kb to ensure no/very low linkage disequilibrium among loci. We chose an admixture model independent of prior population information and with correlated allele frequencies. The algorithm ran with a burnin length of 50’000 and a simulation length of 100’000 Markov chain Monte Carlo (MCMC) repetitions. We varied estimations of *K* between 1 and 10, with 10 repetitions per *K*. The most likely number of populations (*K*) was estimated based on Evanno’s method (Evanno et al. 2005) implemented using the R package *pophelper* version 2.3.0 (Francis 2017). Regions in the genome spanning the most significant associations were further investigated in detail for signatures of linkage disequilibrium. Using the vcftools option --hap-r2, we compared all possible SNP pairs in a 5 kb window. A heatmap was produced based on the *r^2^* values using the R package *LDheatmap* version 0.99-7 (Shin et al. 2006).

### Allelic effect and trade-off analyses

We used *GAPIT* (Tang et al. 2016) for estimations of allelic effects. Allelic effects on EC_50_ values were compared to allelic effects on growth rate and melanization (under temperatures of 18°C, 24°C and 30°C). The total fungicide resistance variation explained by each SNP was determined using a linear mixed-effect model implemented in the *lme4* R package version 1.1-19 (Bates et al. 2015). EC_50_ values were used as response variables, the SNPs as fixed effects and populations were included as random effects. Using the function r.squaredGLMM from the *MuMln* package version 1.43.6 in R we obtained R^2^ indices (Nakagawa & Schielzeth 2013).

### Homology analyses of candidate genes

Amino acid sequences of all genes in the *P. nodorum* SN15 reference genome were obtained from the UniProt database under the proteome ID 000001055 (Magrane & Consortium 2011; Hane et al. 2007). We aligned the predicted protein sequences to the SN2000 genome nucleotide sequences, and incorporated queries with 95% maximum score using the software exonerate version 2.2.0 (Richards et al. 2017; Slater & Birney 2005). For inferences on gene function, we identified conserved domains using InterProScan v.77.0, NCBI Conserved Domain v.3.17 and HMMER v3.3 database search tools (Quevillon et al. 2005; Finn et al. 2011; Marchler-Bauer et al. 2015).

## Results

### Population level differences for fungicide sensitivity and genetic diversity

We analyzed sensitivity to an azole fungicide in a worldwide collection of 159 *P. nodorum* isolates using individual EC_50_ measures. The pathogen strains came from seven field populations located in Australia (*n* = 22), South Africa (*n* = 21), Switzerland 1999A (*n* = 20), Switzerland 1999B (*n* = 22), Iran (*n* = 16), New York (*n* = 21), Oregon (*n* = 16) and Texas (*n* = 21) (Figure 1 A). In total, 39 isolates (24.5%) had EC_50_ values higher than the overall average of 0.12 ppm. The populations from Switzerland showed the highest average EC_50_ values (0.20 and 0.37 ppm in 1999A and 1999B respectively; *P* ≤ 0.01; Figure 1B) while the population from Oregon had the lowest average EC_50_ (0.05 ppm). We sequenced genomes for all 159 isolates using Illumina short-read sequencing. On average, we obtained a mean sequencing depth of 24X per individual, with a SNP density of approximately 12 SNPs per kb. After removing SNPs with more than 10% missing genotypes and minor allele frequencies <5%, we retained a total of 436’365 SNPs to be used for downstream analyses. The total number of SNPs retained per population varied from 340’929 in Australia to 395’054 in Switzerland 1999B. Linkage disequilibrium decayed differently among populations (Figure 2A). In the populations from Australia, Switzerland, Iran and the United States (New York, Oregon, Texas), r^2^ reached values below 0.2 within 10 kb. In the population from South Africa r^2^ ~ 0.2 was reached at 15 kb.

**Fig. 1.**
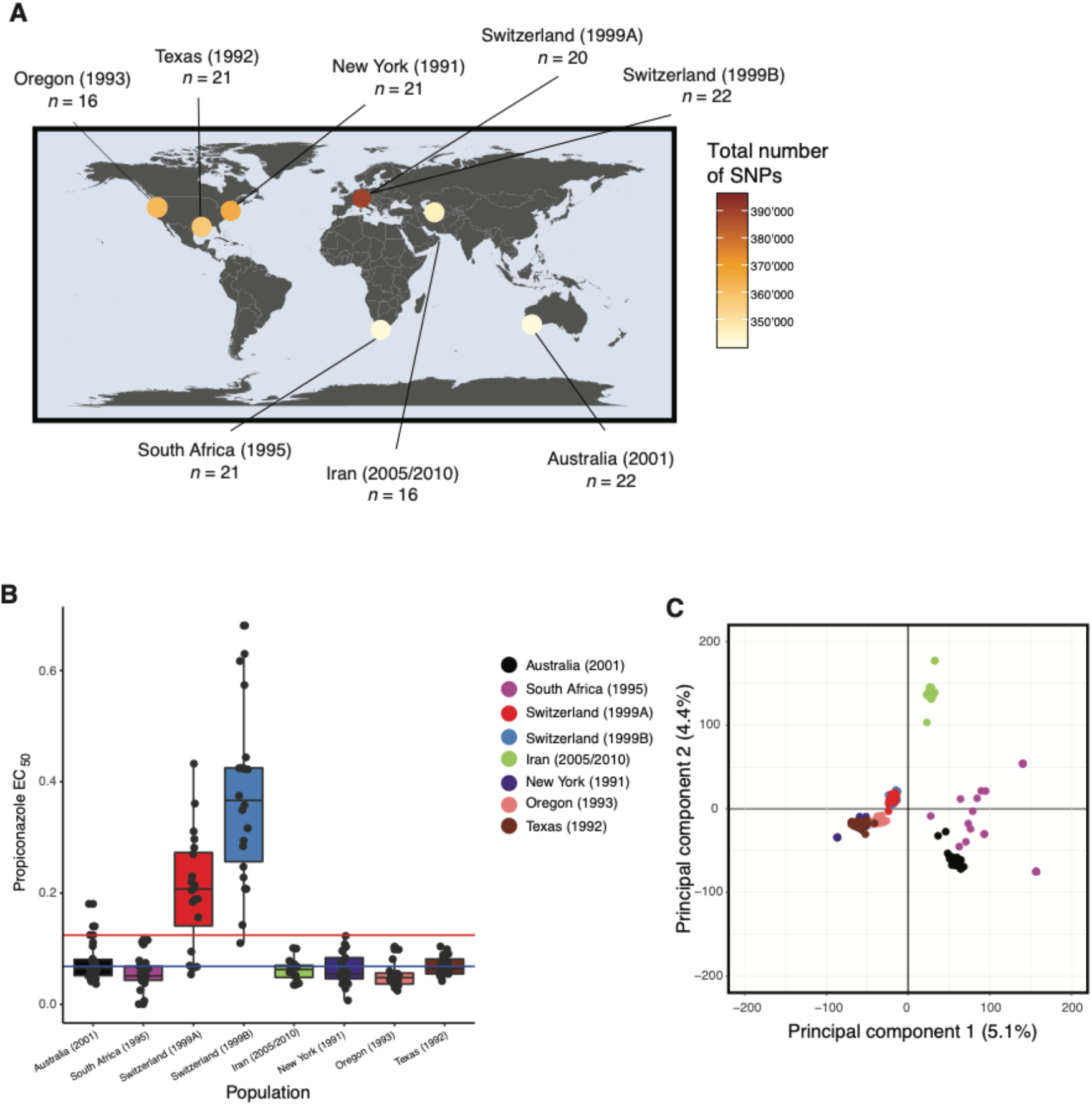
Geographic origins of the 159 *Parastagonospora nodorum* isolates. (*A*) World map showing sampling sites, number of isolates per population, number of single nucleotide polymorphisms, and presence of *CYP51* mutations associated with fungicide resistance. *(B)* Boxplots of EC_50_ (in ppm) values for each isolate in each population. The red line shows the mean overall EC_50_ and the blue line indicates the overall median EC_50_. (*C*) The first two principal components from a PCA of 436’365 SNPs genome-wide SNP genotypes. Populations are color-coded.

A population structure analysis revealed clusters of isolates differentiated according to the continent of origin (Figure 1C). A major cluster was formed by isolates from Switzerland and the United States (New York, Oregon, Texas). Isolates from Australia, South Africa, and Iran constituted a second and more dispersed cluster. We analyzed the clustering scenarios of *K* = 2 and *K* = 3 (Figure S1). At *K* = 2, Australia and South Africa belonged to cluster 1, while the other populations composed cluster 2. At *K*= 3, the Iranian population constituted most of cluster 3 which was shared with genotypes from Switzerland and South Africa (Figure S2). The global dispersal of *P. nodorum* was proposed to mirror the domestication and expansion of the wheat host (Heun 1997; Balter 2007; McDonald et al. 2012; Balfourier et al. 2019). Wheat originated in the Fertile Crescent and then spread across Europe and Asia for thousands of years before Europeans brought it to the American continent ~500 years ago and Australia ~200 years ago. *P. nodorum* is a seedborne pathogen, so it is likely that the pathogen moved globally on infected wheat seed. Because the Iranian *P. nodorum* population is closest to the Fertile Crescent, we expect it would have retained most of the ancestral polymorphism. This is reflected by the finding that it was a hotspot of genetic diversity detected previously by microsatellite markers (McDonald et al. 2012) and neutral SNP markers (Pereira et al. 2019). In Australia, strict quarantine measures likely limited the introduction of the pathogen on infected wheat material (Oliver et al., 2012). Consistent with this proposed bottleneck, we found that the Australian population had low diversity and was distinct from other populations.

### Genetic architecture of fungicide sensitivity across populations

To unravel the genetic architecture of fungicide resistance in *P. nodorum*, we performed genome-wide association analyses using all 159 isolates. We associated genotypes at the 436’365 SNP markers with the EC_50_ phenotypes and identified 34 SNPs significantly associated with fungicide resistance (Table S2). Two associations above the most stringent threshold (Bonferroni a = 0.05, *P* < 1.1 e-07) were located on chromosomes 6 and 15 (Figure 2B). At the FDR 5%, we found five additional associations on chromosomes 12, 15 and 22. At FDR 10%, we identified a total of 27 additional SNPs on chromosomes 2, 4, 7, 8, 9, 10, 15 and 20 (Figure 2B). The average distance between genes in the *P. nodorum* genome is 1.2 kb (Syme et al. 2018). We considered SNPs to be in close proximity if they were located within 1 kb of the closest gene. We found 5 associations > 1 kb from the nearest gene and 16 within 1 kb of a gene. Thirteen associations were located within a gene (Table S2). Quantile-quantile (QQ) plots showed there was not meaningful inflation due to population structure using the MLM + K model (Figure S3).

**Fig. 2.**
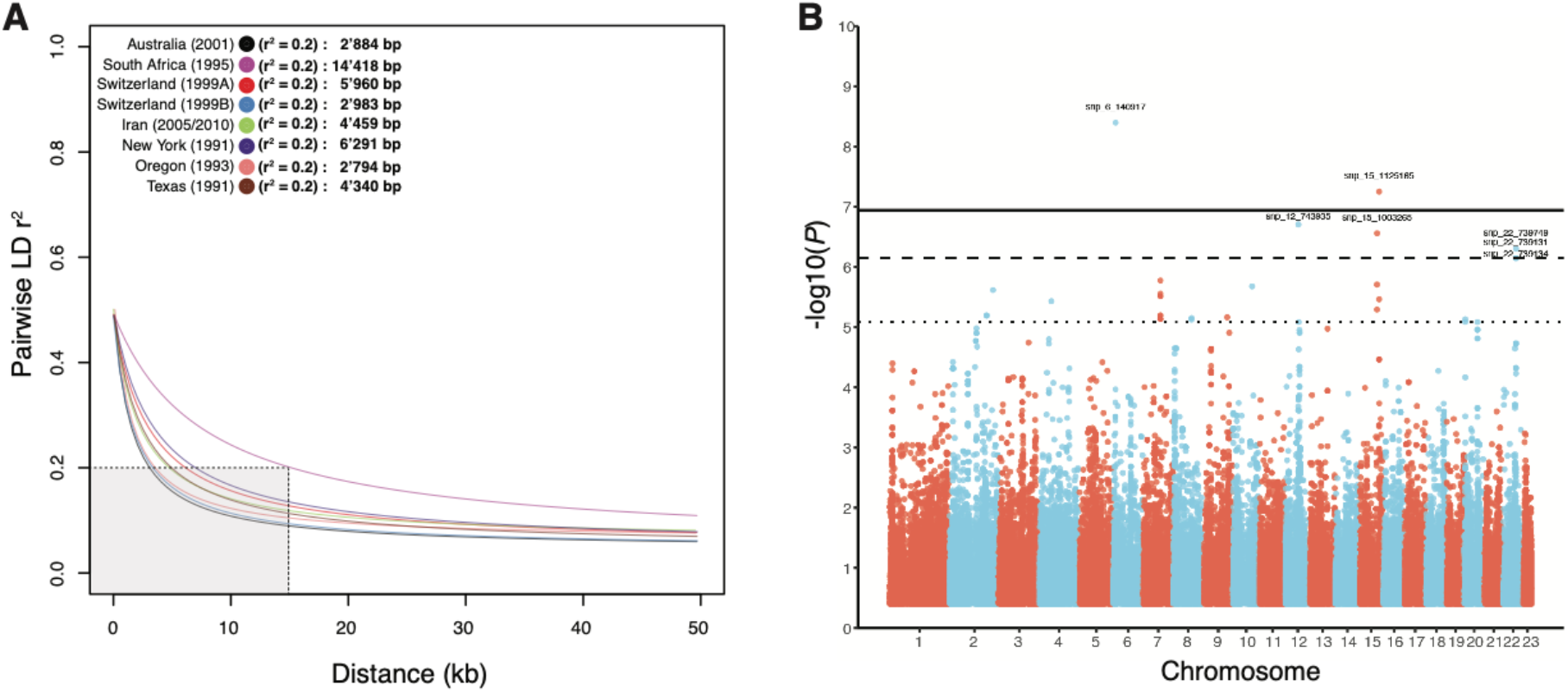
Linkage disequilibrium decay in each population and Manhattan plot of GWAS for fungicide sensitivity. (*A*) Pairwise linkage disequilibrium decay among all SNPs within a fixed window of 50 kb for each population. A non-linear model was fitted based on r^2^ measures along with the first 50 kb on chromosome 1 using the equation of Ingvarsson (2005). The grey shading indicates the total area needed for all populations to reach r^2^ = 0.2. *(B)* Manhattan plot showing the SNP associations with fungicide resistance. SNP markers are shown as dots colored according to their associated chromosomes. Different significance levels are displayed on the y-axis: Horizontal lines represent the thresholds for false discovery rate (FDR) 10% (dotted line), FDR 5% (dashed line) and after Bonferroni correction (solid line). SNPs above FDR 5% were labelled with a specific identifier (chromosome number + SNP coordinates in bp).

Populations differed in their complement of fungicide resistance mutations. Two SNPs above the Bonferroni threshold (snp_6_140917 and snp_15_1125165) were both present in the populations from Switzerland and Texas but were absent in South Africa. The mutation underlying the third strongest association (snp_12_743935, FDR 5%) was exclusively present in the populations from Switzerland but was missing in all other populations. Taken together, based on the top seven SNPs passing the 5% FDR, all populations carried at least one resistance mutation with the exception of South Africa.

### Genomic context of the key loci contributing to azole sensitivity

We investigated the genomic context of the most strongly associated SNPs. The strongest association was snp_6_140917 on chromosome 6 (*P* = 4.03 * 10^-9^, Figure S4). This SNP was located 1044 bp upstream of the nearest gene (SNOG_15057), which encodes a helix-loop-helix (HLH) domain functioning as a transcription factor (Table S2). HLH-domain proteins constitute a large family of proteins acting as gene expression regulators (Massari & Murre 2000). Some members of this family were shown to boost drug resistance gene expression in human tumors (Cheung 2004; Vandeputte et al. 2002) and plant pathogens (Liu et al. 2015). The second most strongly associated genomic region was on chromosome 15 (Figure 3A). The SNPs snp_15_1125165 (*P* = 5.65 * 10^-8^), and snp_15_1124326, (*P* = 3.45 * 10^-6^), were located in close proximity at chromosomal positions 1.125 and 1.124 Mbp, respectively (Figure 3 A). The SNP at 1.125 Mbp comprised a non-synonymous mutation (threonine to isoleucine) in the gene SNOG_14185 (Figure S5). The SNP at 1.124 Mb comprised an intron mutation in the same gene. SNOG_14185 encodes a transmembrane transporter and belongs to the Major Facilitator Superfamily (MFS) with similarity to the Yeast Polyamine transporter 1 (Tpo1). MFS transporters are known multidrug resistance components in model organisms and fungal pathogens (De Rossi et al. 2002). The chromosomal regions surrounding snp_6_140917 and snp_15_1125165 show low linkage disequilibrium (r^2^ < 0.2) with each region harboring only a single gene (Figure S4A, Figure 3A, respectively).

**Fig. 3.**
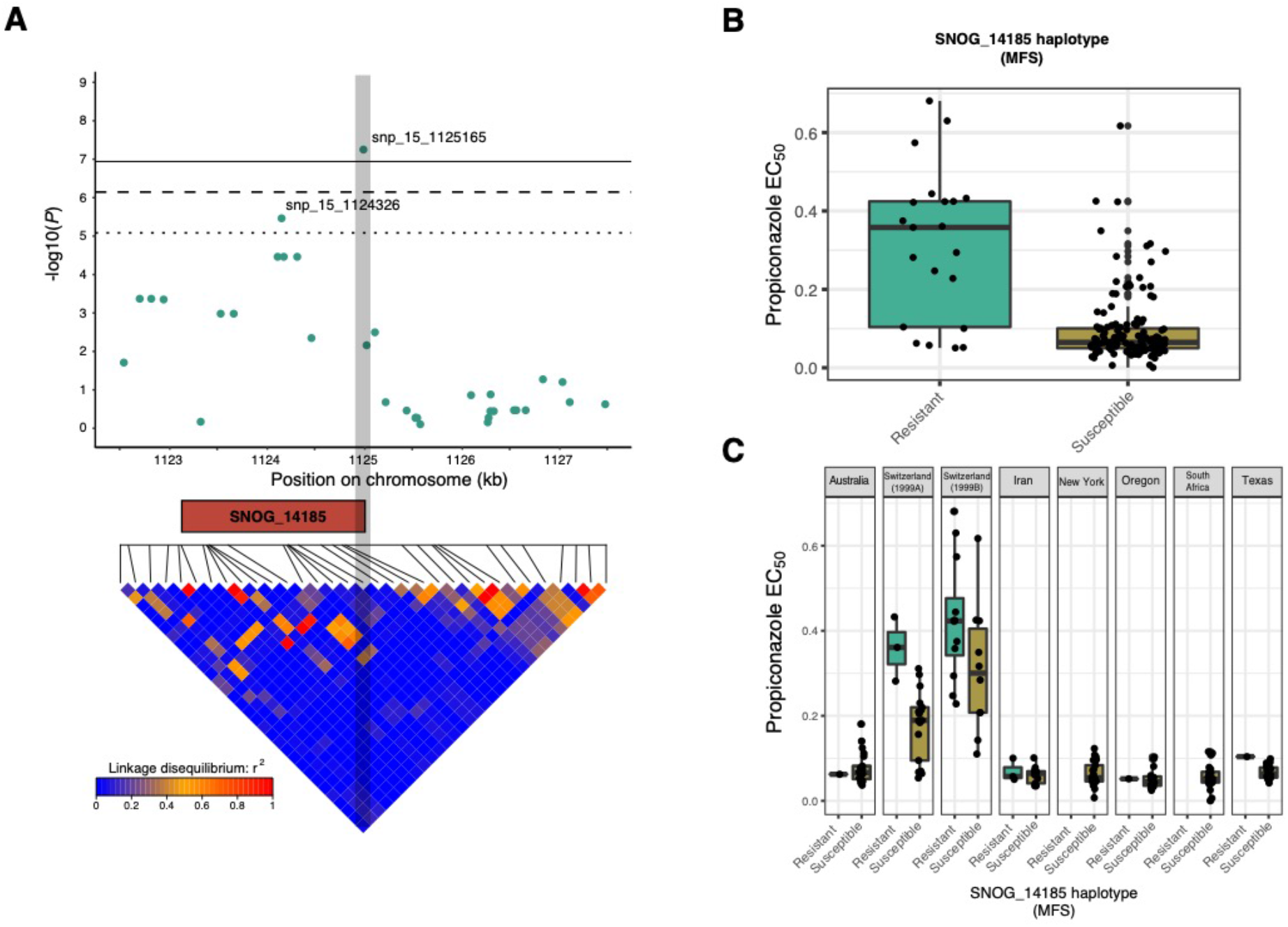
Analysis of the SNP associations near the MFS transporter gene SNOG_14185 associated with azole resistance. (*A*) Top panel: Scatter plot for association *P*-values of SNPs within a 5 kb region centered on the peak snp_15_1125165. Horizontal lines represent the thresholds for false discovery rate (FDR) 10% (dotted line), FDR 5 % (dashed line) and after Bonferroni correction (solid line). SNOG_14185 encoding an MFS transporter is shown in orange. Bottom panel: Linkage disequilibrium (LD) map for the pairwise comparison among SNPs within a 5 kb window. (*B*) Boxplots showing EC_50_ values (in ppm) for propiconazole among global isolates carrying the resistant or susceptible allele at snp_15_1125165. (*C*) Boxplots showing EC_50_ values in isolates carrying the resistant or susceptible allele at snp_15_1125165 organized according to population.

### Combinatorial effects of fungicide resistance loci

We evaluated how the frequencies and effects of the individual SNP associations contributed to the overall azole resistance of *P. nodorum*. The resistance allele at snp_6_140917 was present in 41% of isolates from Switzerland 1999B, in 10% of the New York isolates and in 10% of the Texas isolates (Figure S6 A). The global frequency was about 8% across all 159 isolates. The resistance allele at snp_15_1125165 was present at a global frequency of 14%, at 5% in Australia, 15% in Switzerland 1999A, 55% in Switzerland 1999B, 19% in Iran, 6% in Oregon and 5% in Texas (Figure S6 B). When comparing the degree of fungicide sensitivity, the group of isolates containing either of the two resistance alleles had higher EC_50_ values (Figure 3B, Figure S4). At the population level, the group of isolates harboring the resistant allele at snp_6_140917 was significantly more resistant only within Switzerland 1999B (*t*-test *P* = 0.004; Figure S4). For snp_15_1125165, the group of isolates containing the resistant allele was more resistant within Switzerland 1999A (*t*-test *P* = 0.03) but only marginally more resistant in Switzerland 1999B (*t*-test *P* = 0.08, Figure 3C). In contrast, in the populations from Australia, Iran and Oregon, there were no significant differences between isolates carrying the different alleles.

We expanded the comparisons to genotypes differentiated by non-synonymous mutations in the *CYP51* gene. Isolates with a non-synonymous *CYP51* resistance mutation in the background and a resistance allele identified by GWAS showed a significant increase in resistance in 4 out of 5 combinations (Figure 4). However, when considering only the resistance alleles detected by the GWAS, while disregarding the *CYP51* resistance mutations in the genetic background, we did not find a significant increase in resistance. Next, we assessed the individual contributions of the identified resistance alleles to the overall variation in fungicide sensitivity among populations. The mutations identified in the *CYP51* gene contributed 63.2% of the total phenotypic variation while the snp_15_1125165 in the MFS transporter gene contributed only 6.1% of the phenotype variation.

**Fig. 4.**
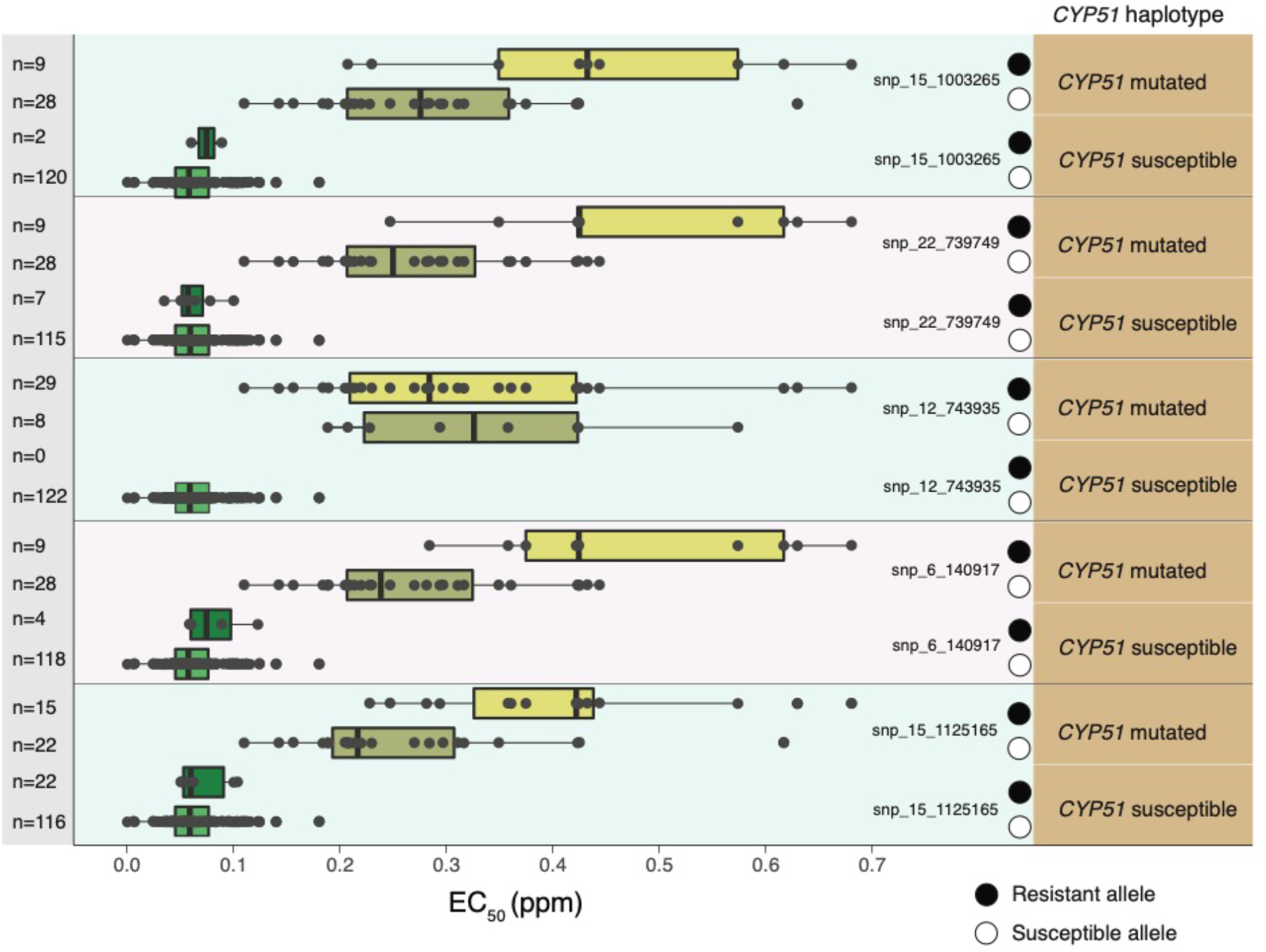
Comparisons of EC_50_ values among resistance haplotypes. Combinations of mutations in the *CYP51* gene and the most significant associations found by GWAS are shown. The number of isolates carrying a specific genotype is indicated on the y-axis.

### Allelic effects associated with fungicide resistance and testing for trade-offs

Allelic effects quantify the mean difference in phenotypic values between genotypes carrying either of two alleles at a locus. We investigated the correlation of allelic effects between fungicide sensitivity and six quantitative life history traits including growth and melanization at the temperatures 18°C, 24°C and 30°C (Figure 5). We focused on the seven most significant SNPs identified in the GWAS for fungicide resistance and performed association mapping analyses for the six other traits. Using allelic effect correlations, we investigated whether resistance mutations showed evidence for pleiotropic effects on any other trait. We found that the most significant SNPs had no meaningful impact on any other analyzed traits (Figures 5A and 5B) and we found no strong correlation between allelic effects of fungicide resistance and the other traits (Figure 5C).

**Fig. 5.**
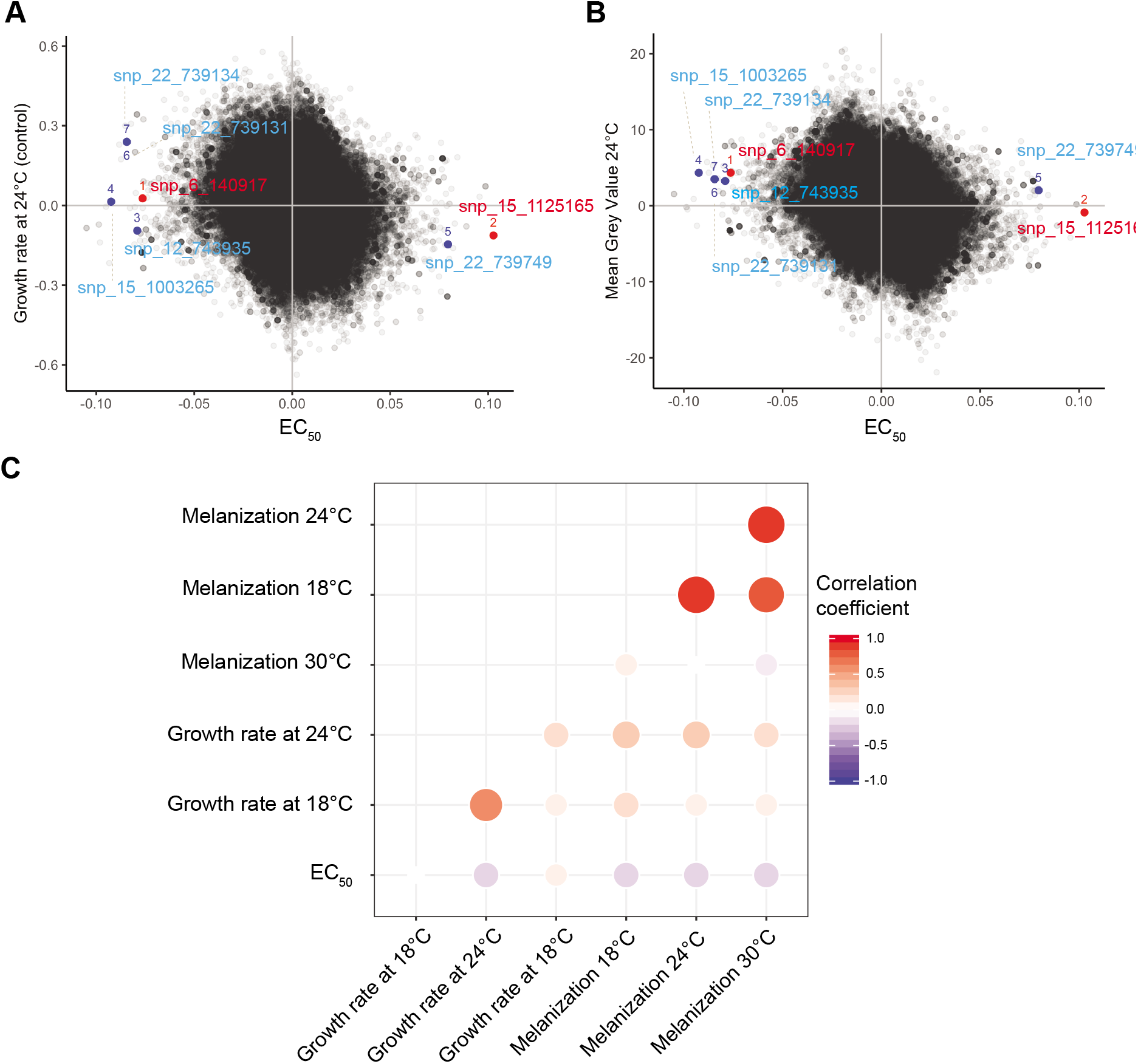
Genome-wide allelic effect correlations. Allelic effects for EC_50_ and *(A)* growth rate and (*B*) melanization at 24°C. SNPs that were significantly associated with propiconazole sensitivity are indicated. (*C*) Mean allelic effect correlation coefficients for EC_50_, growth rate (mm day^-1^) and melanization at different temperatures. Sizes of circles represent degrees of significance and the opacity of colors are proportional to the size of the correlation coefficient.

## Discussion

We used whole-genome sequencing data and association mapping to reveal the multilocus genetic architecture of fungicide resistance in *P. nodorum*. We identified significant differences in azole resistance among a global set of field populations. Some populations evolved distinct combinations of resistance alleles which showed synergistic interactions. We identified several significantly associated SNPs in close proximity to candidate resistance genes, including an MFS transporter. There was no evidence for trade-offs associated with the observed resistance to azoles.

The genetic basis of fungicide resistance includes both qualitative and quantitative factors (Waard et al. 2006). The presence or absence of a sensitive target site is typically considered a qualitative factor (e.g, *Strobilurus tenacellus* and strobilurin A (Kraiczy et al. 1996)). Previous studies oriented around single known loci identified major genetic determinants (e.g. qualitative factors) associated with fungicide resistance in *P. nodorum* (Blixt et al. 2009; Pereira et al. 2017). Quantitative factors are often associated with a number of different mechanisms that make minor contributions to overall resistance. In this study we used a genome-wide approach to identify both major and minor contributions to resistance. We found 34 candidate loci distributed across the genome, including the *CYP51* gene, underlying quantitative variation in fungicide sensitivity across populations. In clinical resistance studies, an increasing number of genetic loci affecting drug resistance have been described in viruses, protozoa, and bacteria (Manolio 2013; Chewapreecha et al. 2014; Holt et al. 2015; Power et al. 2017). The emergence of fungicide resistance in plant pathogenic fungi has been associated mainly with mutations in genes encoding the targeted protein. However, GWAS based on whole-genome sequencing in the barley scald pathogen *Rhynchosporium commune* (Mohd-Assaad et al. 2016) showed that *R. commune* evolved resistance to azoles via a combination of genetic variants in addition to mutations in the *CYP51* gene.

The evolution of azole resistance in *P. nodorum* was likely initiated by mutations in the *CYP51* gene coupled with more recent mutations in other loci. In both *P. nodorum* and *R. commune*, the mutations with the greatest impact on azole resistance were found in the *CYP51* gene (Mohd-Assaad et al. 2016). The convergent evolution of azole resistance based on *CYP51* mutations is a major feature of azole resistance globally (Fisher et al. 2018). However, additional loci may elevate fungicide resistance in a subset of populations. There is growing evidence that herbicide resistance in plants involves sites that are not targeted by the herbicide (Baucom 2019). These non-target sites are usually related to herbicide translocation or detoxification (Peng et al. 2010; Leslie & Baucom 2014).

We observed different levels of resistance and combinations of resistance alleles among our worldwide populations. Such a mosaic structure in resistance factors was also observed among populations of *Streptococcus pneumoniae* and *Mycobacterium tuberculosis* (Chewapreecha et al. 2014; Farhat et al. 2019), with certain populations enriched in particular resistance determinants (Chewapreecha et al. 2014). The highest number of resistance alleles was found in *P. nodorum* isolates from Switzerland (both from 1999A and 1999B), which were also the most resistant populations. The use of azoles in Europe started in 1979 (Estep et al. 2015). We postulate that the Swiss populations were either directly selected for fungicide resistance or received resistance alleles through gene flow. Considering migration patterns based on microsatellite markers (Stukenbrock et al. 2006), it is likely that resistance genotypes will be exchanged among populations. Populations of the barley pathogen *R. commune* showed a similar pattern, with the Swiss population among the most resistant worldwide and an exchange of migrants among continents (McDonald 2015; Mohd-Assaad et al. 2016). Populations of the wheat pathogen *Zymoseptoria tritici* from Oregon (USA) acquired fungicide resistance over a period of two decades (Estep et al. 2015). Given that we analyzed European populations many years after the onset of widespread fungicide applications, many mutations affecting azole resistance may have arisen in these populations.

We identified a potentially new mechanism of azole resistance in *P. nodorum*. Isolates with the lowest sensitivity to propiconazole often harbored resistance mutations at both the *CYP51* locus and the MFS transporter locus. MFS transporters are among the largest protein families (Stergiopoulos et al. 2002), ubiquitous in the cell membrane of prokaryotes (Hirai & Subramaniam 2004) and eukaryotes (Henderson & Maiden 1990). These transporters contribute to cell-to-cell communication as well as movement of pathogenicity toxins and antimicrobial drugs through the cell membrane (Paulsen et al. 1996). Importantly, MFS transporters can also act as efflux pumps that reduce intracellular drug concentrations (Kretschmer et al. 2009; Omrane et al. 2015; Redhu et al. 2016). In the plant pathogens *Botrytis cinerea and Z. tritici*, upregulation of an MFS transporter was shown to reduce sensitivity to azole fungicides (Kretschmer et al. 2009; Omrane et al. 2017). In *P. nodorum*, we observed correlations between MFS mutations and azole sensitivity. MFS transporters can also vary in their substrate affinity as found for a multidrug transporter in *C. albicans* (Pasrija et al. 2007). Depending on the type of MFS transporter mutations, *C. albicans* varied in sensitivity to different drugs, including an azole. The non-synonymous mutation we identified in *P. nodorum* could influence this MFS transporter’s affinity for propiconazole. Interestingly, the group of isolates lacking *CYP51* resistance mutations, but carrying the more resistant variant of the MFS transporter were highly susceptible, indicating that the MFS transporter mutations depend on *CYP51* mutations to have an effect. This is similar to what was observed in *Z. tritici* (Omrane et al. 2015). Epistasis among resistance-encoding genes is also known from *C. albicans* (Hill et al. 2013; Ciudad et al. 2016) and appears to be a common phenomenon associated with the emergence of *de novo* resistance mutations.

A major constraint on the emergence of resistance mutations is negative pleiotropy. Fungicides generally target essential metabolic processes. By reducing the synthesis of ergosterol, azoles negatively impact cell fluidity and functions through membrane defects (Georgopapadakou & Walsh 1996; Lass-Flörl 2011). Resistance mutations are most successful if they confer decreased binding affinity with the fungicide but do not negatively impair normal protein functions (Karaoglanidis et al. 2001; Yan et al. 2011). Resistance mutations that lead to over-production of targeted proteins may negatively impact the cellular energy budget (Lang et al. 2009). Hence, resistance mutations are likely to confer advantageous effects only in the presence of the fungicide. Fitness costs in the absence of the pesticide constrain the emergence of acquired resistance in plants, bacteria, and fungi (Schenk & de Visser 2013; Moura de Sousa et al. 2017; Pagnout et al. 2019). Interestingly, we found no evidence that the most important fungicide resistance mutations negatively impacted growth rates or melanization in *P. nodorum*. This is in contrast to other fungal pathogens such as *Z. tritici* and *R. commune* where growth rates were negatively affected in the absence of azoles (Lendenmann et al. 2015; Mohd-Assaad et al. 2016). Herbicide resistance shows a broader spectrum of pleiotropic effects among species (Powles & Yu 2010; Baucom 2019) with the most common mutations having limited fitness costs (Tranel & Wright 2002). Fitness costs of resistance mutations can also be reduced through compensatory mutations, as shown in bacteria (Levin et al. 2000; Trindade et al. 2009; Moura de Sousa et al. 2017) and postulated in fungi (Lucas et al. 2015; Dettman et al. 2017). We also found no evidence for genetic trade-offs between fungicide resistance and growth at different temperatures.

These findings suggest that *P. nodorum* either evolved azole resistance without relying on costly mutations that would affect other traits or that trade-offs have already been resolved through fixed compensatory mutations. The negative pleiotropic effects associated with resistance mutations can be masked by compensatory mutations occurring in the genetic background (Becher & Wirsel 2012; Cools et al. 2013). Fitness costs could also manifest for different traits than we analyzed (e.g. virulence, competitive ability, etc.) or depend on specific environmental conditions that we did not consider.

Our study demonstrates how genome-wide association studies of a global collection of pathogen strains can recapitulate the emergence of fungicide resistance. The distinctive complements of resistance mutations found among populations reflects how the evolutionary trajectory of fungicide adaptation is complex and difficult to predict. The apparent lack of tradeo-ffs to adapt to azole fungicides in *P. nodorum* highlights how more sustainable crop protection strategies are needed. An absence of trade-offs will contribute to a rapid decline in fungicide effectiveness and more widespread losses in crop production.

## Acknowledgments

We thank Marcello Zala for assistance in the laboratory and Lea Stauber for helpful comments on a previous version of this manuscript. The Genetic Diversity Center (GDC) of ETH Zurich and the Functional Genomics Center in Zurich provided sequencing facilities. This study was financed in part by the Coordenação de Aperfeiçoamento de Pessoal de Nível Superior - Brasil (CAPES) - Finance Code 001.

