## Supplementary Information for "The genetic architecture of emerging fungicide resistance in populations of a global wheat pathogen"

### Supplementary Figures

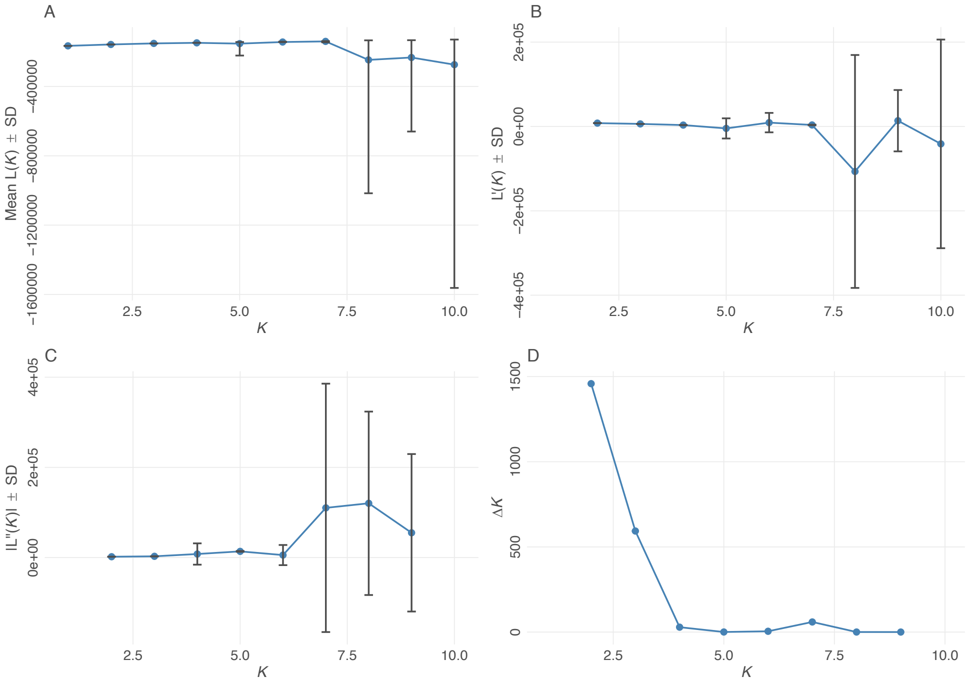


**Supplementary Figure 1: Evaluation of the cluster assignments using STRUCTURE.** The panels show the mean estimated *ln* probability of the data, based on the first derivative per K and delta K. (A) Mean likelihood and variance per *K* value. (B) The rate of change of the likelihood distribution. (C) The absolute value of the second order rate of change of the likelihood distribution. (D) Mean delta *K* plot from K=2 to 9.

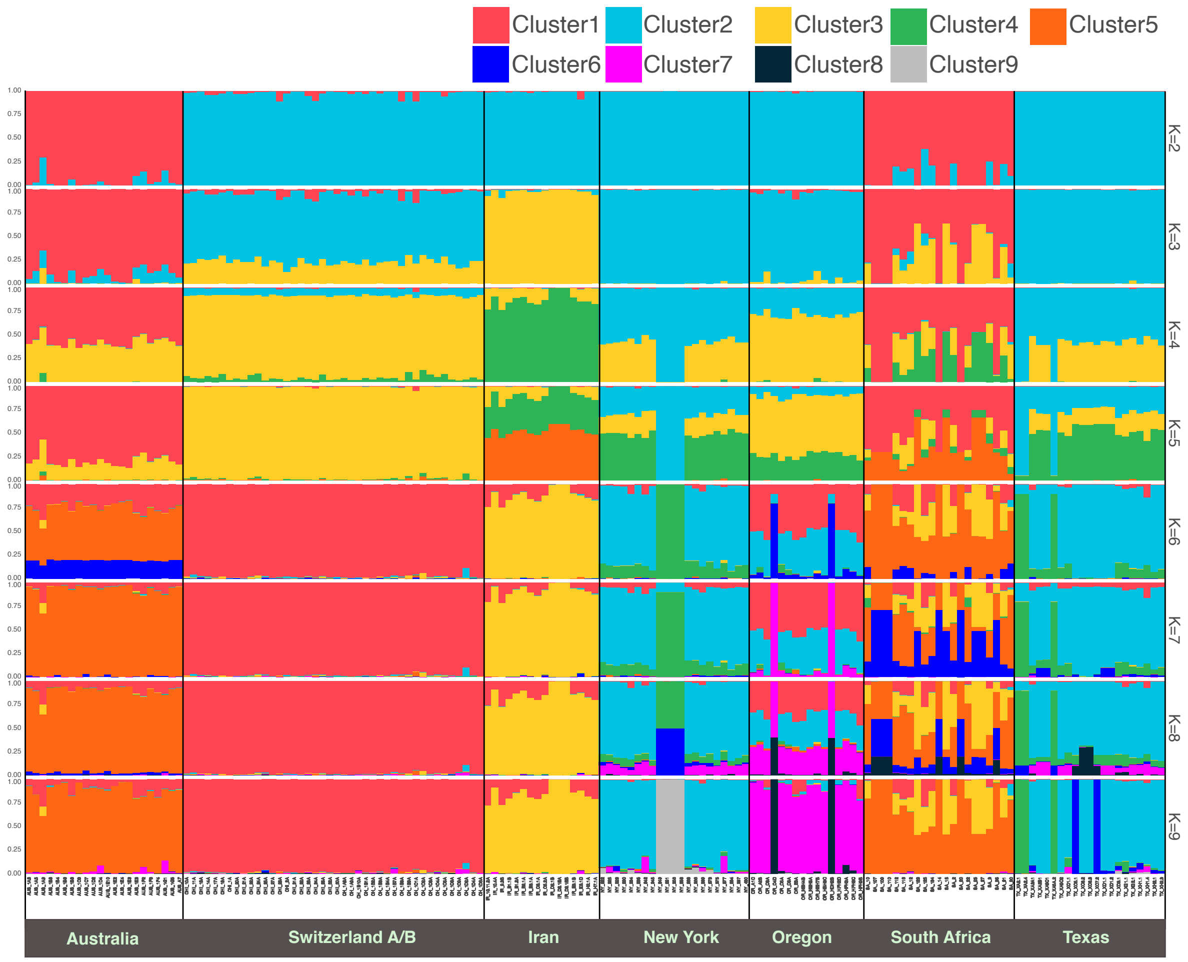


**Supplementary Figure 2. Bayesian genetic clustering of 159 isolates of *Parastagonospora nodorum* based on 2348 single nucleotide polymorphic markers using STRUCTURE.** Vertical colored bars represent the assignment probability of each isolate for different values of K.

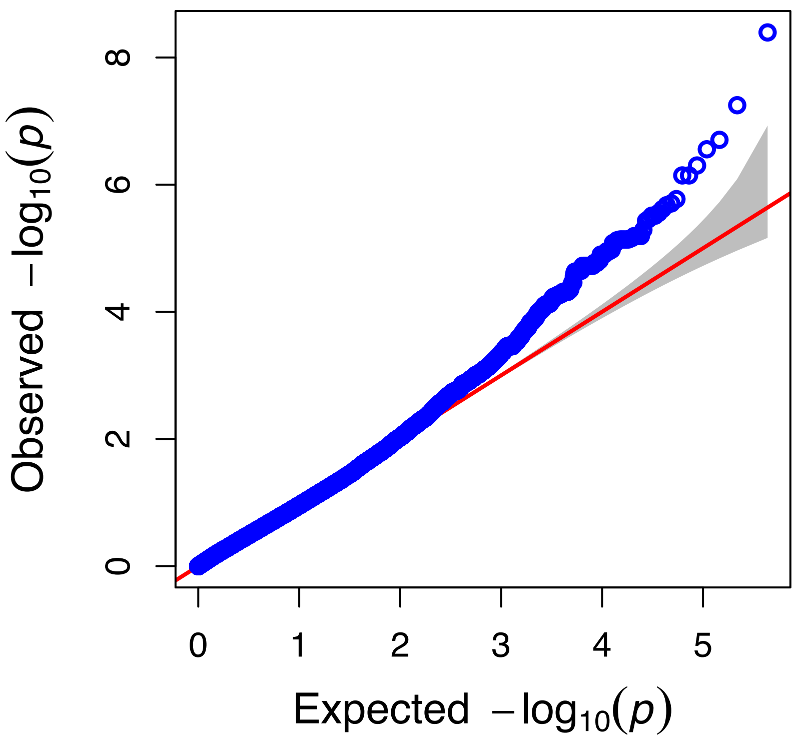


**Supplementary Figure 3: Quantile-Quantile (QQ) plots.** The -$\log_{10}$ *P*-values obtained for the genome wide association using $EC_{50}$ scores (blue dots). The red continuous line indicates the expected values and the grey interval shows the confidence interval.

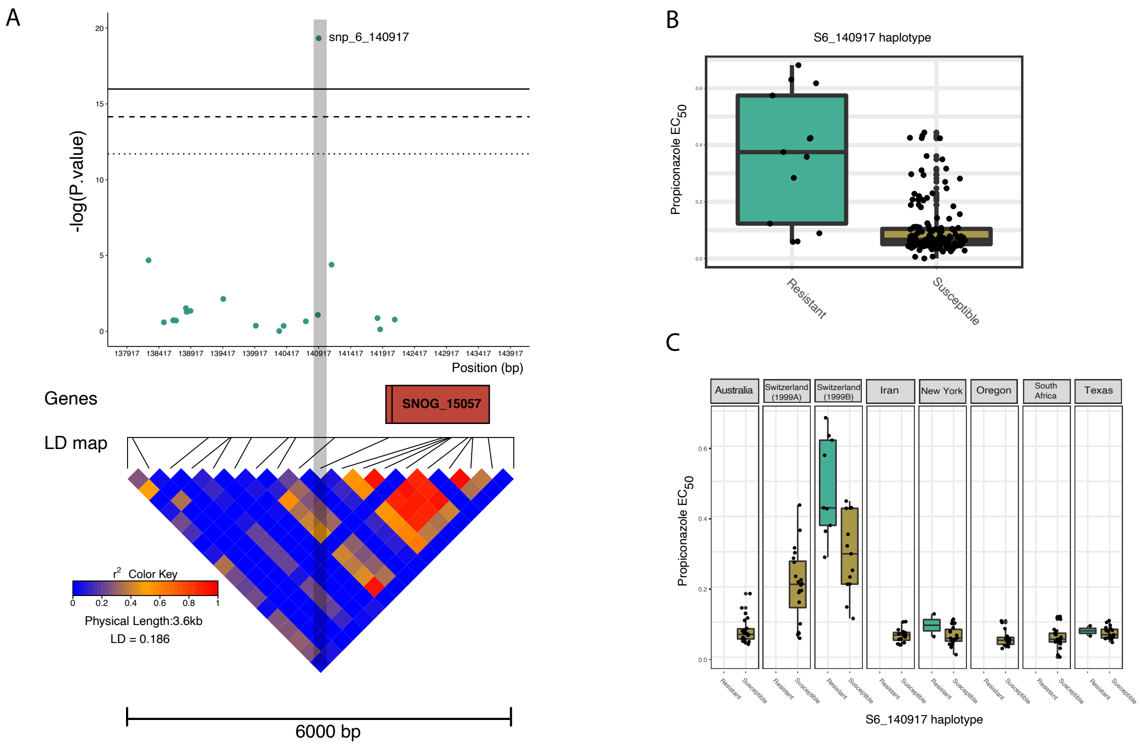


**Supplementary Figure 4**: **Detailed analysis of candidate markers associated with fungicide sensitivity in *Parastagonospora nodorum*.** (A) *P*-value distribution around snp_6_140917. SNOG_15057 encoding a helix-loop-helix (HLH) domain is shown in orange. The heatmap of pairwise linkage disequilibrium $r^2$ for SNPs considering all 159 isolates. The region spans a total of 6 kbp. (B) Boxplots of $EC_{50}$ values for the group of isolates possessing the resistant versus the susceptible allele at snp_6_140917. (C) Boxplots of $EC_{50}$ values between isolates carrying the resistant versus the susceptible allele organized according to population.

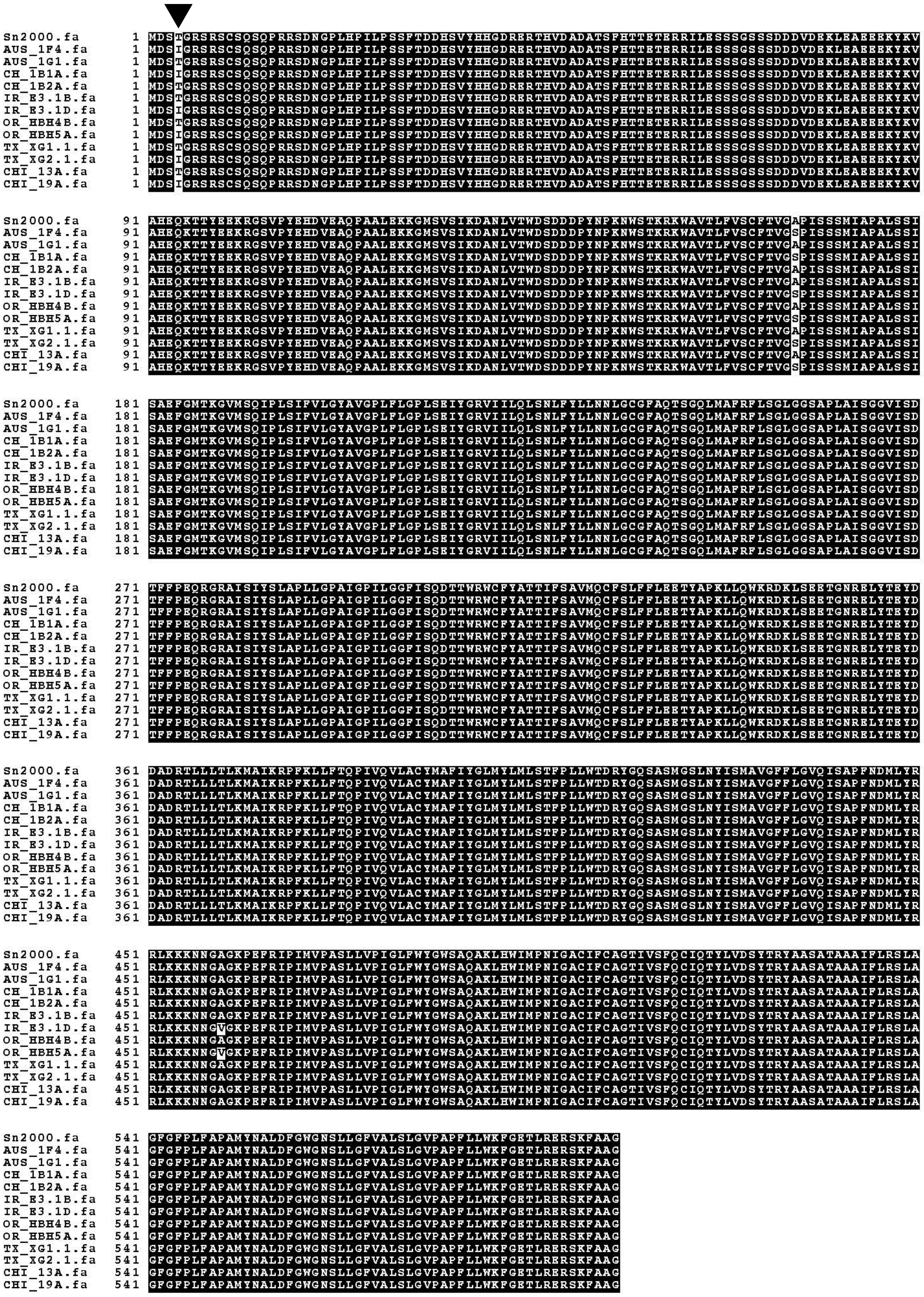

**Supplementary Figure 5**: Multiple sequence alignment for the SNOG_14185 amino acid sequence showing isolates from the global distribution. The black triangle in the amino acid position four indicate the non-synonymous change identified in the GWAS. Isolates harboring an isoleucine (I) were found to be associated to higher EC50 values compared to isolates containing a threonine (T).

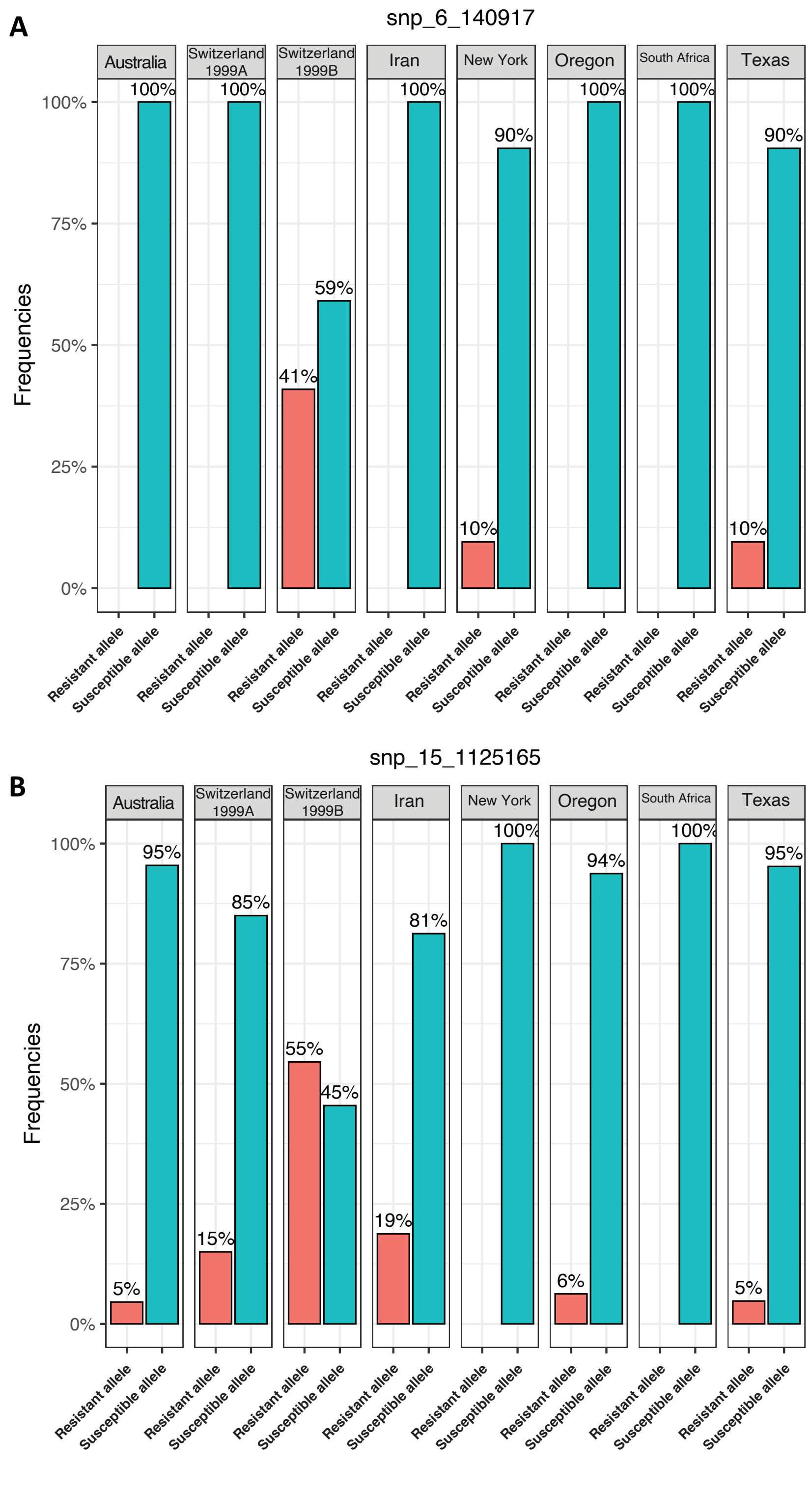


**Supplementary Figure 6**: Frequencies of resistant and susceptible alleles within populations of *Parastagonospora nodorum*. (*A*) SNP maker snp_6_140917 and (*B*) snp_15_1125165.

### Supplementary Tables

**Supplementary Table 1. Number of principal components (PCs) included in the GWAS, the Bayesian Information Criteria (BIC) value and the likelihood values.**

| Number of PCs/Covariates | BIC (larger is better) - Schwarz 1978 | log Likelihood Function Value |
|---|---|---|
| 0 | 155.23 | 162.83 |
| 1 | 153.04 | 163.18 |
| 2 | 150.60 | 163.28 |
| 3 | 150.50 | 165.70 |

**Supplementary Table 2. The top loci from the genome-wide association mapping in *Parastagonospora nodorum* for different significance thresholds. Coordinates and distances are given in base pairs.**

| Threshold | p-value | SNP | Chromosome | Position | P.value | maf | Start_gene | End_gene | gene ID | distance |
|---|---|---|---|---|---|---|---|---|---|---|
| Bonferonni | 1.16E-07 | S6_140917 | 6 | 140917 | 4.03E-09 | 0.08 | 141961 | 143580 | SNOG_15057 | -1044 |
| Bonferonni | 1.16E-07 | S15_1125165 | 15 | 1125165 | 5.62E-08 | 0.14 | 1123292 | 1125175 | SNOG_14185 | 0 |
| FDR_5% | 7.15E-07 | S12_743935 | 12 | 743935 | 1.97E-07 | 0.19 | 742669 | 743805 | SNOG_03692 | 130 |
| FDR_5% | 7.15E-07 | S15_1003265 | 15 | 1003265 | 2.77E-07 | 0.08 | 1001315 | 1002569 | SNOG_14240 | 696 |
| FDR_5% | 7.15E-07 | S22_739749 | 22 | 739749 | 4.99E-07 | 0.12 | 739857 | 741023 | SNOG_10460 | -108 |
| FDR_5% | 7.15E-07 | S22_739131 | 22 | 739131 | 7.15E-07 | 0.09 | 738155 | 738916 | SNOG_10462 | 215 |
| FDR_5% | 7.15E-07 | S22_739134 | 22 | 739134 | 7.15E-07 | 0.09 | 738155 | 738916 | SNOG_10462 | 218 |
| FDR_10% | 8.26E-06 | S7_999491 | 7 | 999491 | 1.69E-06 | 0.16 | 999693 | 1002904 | SNOG_06551 | -202 |
| FDR_10% | 8.26E-06 | S15_995131 | 15 | 995131 | 1.97E-06 | 0.12 | 993566 | 994895 | SNOG_14244 | 236 |
| FDR_10% | 8.26E-06 | S10_1112832 | 10 | 1112832 | 2.10E-06 | 0.06 | 1111907 | 1112980 | SNOG_05992 | 0 |
| FDR_10% | 8.26E-06 | S2_2489215 | 2 | 2489215 | 2.42E-06 | 0.06 | 2486059 | 2486877 | SNOG_30129 | 2338 |
| FDR_10% | 8.26E-06 | S7_1000371 | 7 | 1000371 | 2.78E-06 | 0.14 | 999693 | 1002904 | SNOG_06551 | 0 |
| FDR_10% | 8.26E-06 | S7_999855 | 7 | 999855 | 3.07E-06 | 0.14 | 999693 | 1002904 | SNOG_06551 | 0 |
| FDR_10% | 8.26E-06 | S7_999973 | 7 | 999973 | 3.07E-06 | 0.14 | 999693 | 1002904 | SNOG_06551 | 0 |
| FDR_10% | 8.26E-06 | S15_1124326 | 15 | 1124326 | 3.45E-06 | 0.12 | 1123292 | 1125175 | SNOG_14185 | 0 |
| FDR_10% | 8.26E-06 | S4_690420 | 4 | 690420 | 3.72E-06 | 0.24 | 688597 | 689699 | SNOG_30188 | 721 |
| FDR_10% | 8.26E-06 | S15_997120 | 15 | 997120 | 5.12E-06 | 0.13 | 995750 | 996587 | SNOG_14243 | 533 |
| FDR_10% | 8.26E-06 | S2_2115190 | 2 | 2115190 | 6.43E-06 | 0.06 | 2113412 | 2114494 | SNOG_02160 | 696 |
| FDR_10% | 8.26E-06 | S2_2115259 | 2 | 2115259 | 6.43E-06 | 0.06 | 2116921 | 2118744 | SNOG_02157 | -1662 |
| FDR_10% | 8.26E-06 | S7_1000797 | 7 | 1000797 | 6.44E-06 | 0.12 | 999693 | 1002904 | SNOG_06551 | 0 |
| FDR_10% | 8.26E-06 | S7_999559 | 7 | 999559 | 6.44E-06 | 0.12 | 999693 | 1002904 | SNOG_06551 | -134 |
| FDR_10% | 8.26E-06 | S9_1398308 | 9 | 1398308 | 6.86E-06 | 0.07 | 1398685 | 1405487 | SNOG_08614 | -377 |
| FDR_10% | 8.26E-06 | S8_1059071 | 8 | 1059071 | 7.07E-06 | 0.06 | 1059052 | 1059496 | SNOG_07538 | 0 |
| FDR_10% | 8.26E-06 | S7_999266 | 7 | 999266 | 7.28E-06 | 0.12 | 998455 | 999165 | SNOG_30471 | 101 |
| FDR_10% | 8.26E-06 | S7_999349 | 7 | 999349 | 7.28E-06 | 0.12 | 998455 | 999165 | SNOG_30471 | 184 |
| FDR_10% | 8.26E-06 | S7_999820 | 7 | 999820 | 7.28E-06 | 0.12 | 999693 | 1002904 | SNOG_06551 | 0 |
| FDR_10% | 8.26E-06 | S7_999823 | 7 | 999823 | 7.28E-06 | 0.12 | 999693 | 1002904 | SNOG_06551 | 0 |

| | | | | | | | | | | |
|---|---|---|---|---|---|---|---|---|---|---|
| FDR_10% | 8.26E-06 | S7_999307 | 7 | 999307 | 7.28E-06 | 0.12 | 998455 | 999165 | SNOG_30471 | 142 |
| FDR_10% | 8.26E-06 | S7_999816 | 7 | 999816 | 7.28E-06 | 0.12 | 999693 | 1002904 | SNOG_06551 | 0 |
| FDR_10% | 8.26E-06 | S20_60249 | 20 | 60249 | 7.42E-06 | 0.18 | 61041 | 63647 | SNOG_12292 | -792 |
| FDR_10% | 8.26E-06 | S8_1057002 | 8 | 1057002 | 7.50E-06 | 0.06 | 1054678 | 1057518 | SNOG_07536 | 0 |
| FDR_10% | 8.26E-06 | S20_60214 | 20 | 60214 | 8.21E-06 | 0.14 | 56340 | 58034 | SNOG_12290 | 2180 |
| FDR_10% | 8.26E-06 | S20_763661 | 20 | 763661 | 8.26E-06 | 0.11 | 767389 | 768302 | SNOG_13193 | -3728 |
| FDR_10% | 8.26E-06 | S12_763273 | 12 | 763273 | 8.26E-06 | 0.16 | 762821 | 764490 | SNOG_03702 | 0 |
